# Red-beet betalain pigments inhibit amyloid-β aggregation and toxicity in amyloid-β expressing *Caenorhabditis elegans*

**DOI:** 10.1101/2020.12.23.424246

**Authors:** Tomohiro Imamura, Hironori Koga, Yasuki Higashimura, Noriyoshi Isozumi, Kenji Matsumoto, Shinya Ohki, Masashi Mori

## Abstract

**BACKGROUND:** Betalain pigments are mainly produced by plants in the order Caryophyllales. Recent interest in the biological functions of betalain pigments has increased with antioxidant, anti-inflammatory, and anticancer activities reported.

**RESULTS:** We investigated the effects of betalain pigments derived from red-beet on amyloid-β (Aβ) aggregation, a cause of Alzheimer’s disease. Inhibition of Aβ aggregation against Aβ40 and Aβ42 by betalain pigments *in vitro* was demonstrated by the Thioflavin T fluorescence assay, circular dichroism spectroscopy analysis and transmission electron microscopic observations. Moreover, we examined the ability of betalain pigments to interfere with Aβ toxicity by using the transgenic *Caenorhabditis elegans* strain CL2006, which expresses the human Aβ42 protein intracellularly within the body wall muscle and responds to Aβ-toxicity with paralysis. Treatment with 50 μM betalain pigments significantly delayed the paralysis of *Caenorhabditis elegans*.

**CONCLUSION:** These results suggest that betalain pigments reduce Aβ-induced toxicity by inhibiting Aβ aggregation and may lead to their use as inhibitors of Aβ aggregation.

## INTRODUCTION

Betalains are tyrosine-derived pigments found exclusively in plants of the order Caryophyllales,^1^ some *Basidiomycota* fungi^2^ and one bacterial species, *Gluconacetobacter diazotrophicus*.^3^ Betalain pigments are divided into betacyanins (red and purple) and betaxanthins (yellow and orange). Unlike other plant pigments, e.g., flavonoids and carotenoids, betalains contain nitrogen. Previous studies have demonstrated that betalains exhibit strong antioxidant activity^4^ and are involved in plant responses to environmental stimuli and stress.^5, 6^ Betalains also have commercial applications; they are often used as food additives because of their vivid colors. For example, beetroot extract, designated by the label E162, is approved by the United States Food and Drug Administration (FDA) and European Union regulatory agencies for use as a natural food colorant.

Biological activities of betalain pigments have been reported. Betalain-rich extracts possess anti-inflammatory, anticancer and anti-diabetic properties.^7–10^ Furthermore, betanin, the major red pigment of beetroot extract, has been demonstrated to inhibit low-density lipoprotein oxidation.^11, 12^ Our recent study found that the amaranthin betalain pigment, which accumulates in amaranth and quinoa, inhibits HIV-1 protease activity.^13^ Although various physiological functions of the betalain pigments have been reported, it is envisaged that many of the physiological activities of these pigments remain undiscovered.

Alzheimer’s disease (AD) is an irreversible, progressive brain disorder that progressively impairs memory and the ability to think.^14^ From a neuropathological viewpoint, AD is characterized by the aggregation of the amyloid-β (Aβ) protein in senile plaques and hyperphosphorylated tau protein in brain neurofibrillary tangles.^15^ Thus, inhibiting the aggregation and accumulation of Aβ and phosphorylated tau in the brain is the key to treating AD. However, all current FDA-approved anti-AD drugs are only symptomatic treatments, which do not cure the disease. Recent studies have reported the efficacy of plant-derived polyphenolic compounds as candidate substances for AD treatment.^16, 17^ Polyphenolic compounds such as resveratrol, curcumin, quercetin, genistein, morin, and epigallocatechin gallate have been shown to inhibit Aβ production and aggregation.^18, 19^

*Caenorhabditis elegans* (*C. elegans*) is currently used in many studies as a model organism for AD.^20^ *C. elegans* is a free-living, non-parasitic nematode. Approximately 38% of the *C. elegans* genome has human orthologs, including AD-related orthologs such as the *amyloid-bata precursor protein* and *tau*.^21^ Therefore, *C. elegans* has many advantages as an *in vivo* model for AD research. Moreover, a transformant strain CL2006 expressing the human amyloid gene has been generated.^22^ This transformed strain exhibits a paralytic phenotype by accumulating human amyloid-β 1–42 (Aβ42) in muscle.^22^ This transformant phenotype is useful for evaluating substances that are effective against AD. Many studies using this nematode to examine the bioactivity of phytochemicals in Aβ aggregation have been reported;^23–25^ however, the bioactivity of betalain pigments have not been reported in AD studies using nematodes.

In this study, we evaluated the biological activity of betalain pigments against Aβ aggregation using red-beet betalain pigments, mainly composed of betanin and isobetanin. The *in vitro* ThT fluorescence assay and circular dichroic (CD) spectroscopy analysis were performed to evaluate the effect of red-beet betalain pigments on Aβ aggregation. Furthermore, sample of Aβ with and without treatment with red-beet betalain pigments were examined by transmission electron microscopy (TEM). In all experiments, red-beet betalain pigments were found to inhibit Aβ aggregation *in vitro*. Additionally, we evaluated the biological activity of red-beet betalain pigments using an Aβ expressing nematode strain CL2006. Feeding red-beet betalain pigments to Aβ expressing *C. elegans* alleviated the severity of the phenotype associated with Aβ aggregation.

## MATERIAL AND METHODS

### Materials

Lyophilized human amyloid-β 1–40 (Aβ40) and amyloid-β 1–42 (Aβ42) were obtained from SCRUM Inc. (Tokyo, Japan). Red-beet betalains (red-beet betanin extract diluted with dextrin) were obtained from Tokyo Chemical Industry (TCI, Tokyo, Japan). Morin hydrate (TCI) was used as a control for the ThT fluorescence assay. Dextrin (TCI) was used as a control in nematode experiments.

### Sample preparation

Red-beet betalain pigments were mixed with dextrin for preservation. Betalain pigments were purified from red-beet betalain pigments to remove any side effects of dextrin when conducting *in vitro* experiments. First, an aqueous solution of red-beet betalain pigment was prepared and added to DEAE Sephacel (Pharmacia, Uppsala, Sweden). The solution was washed with water before eluting the pigments with 300 mM NaCl. Next, the eluate was adsorbed on cellulose powder (Fuji Film, Osaka, Japan) and washed three times with ethanol. Subsequently, the pigments were eluted with water and concentrated. The eluate was added to reversed-phase Cosmosil 140C_18_-OPN (Nacalai tesque, Kyoto, Japan), washed with water and eluted with 10% acetonitrile. The eluate was evaporated to dryness and residues were dissolved in water and stored at −20 °C until needed. These purified red-beet betalain pigments were used for *in vitro* experiments.

### Preparation of Aβ monomer

Aβ monomer was prepared according to previous studies.^26, 27^ Aβ was dissolved in 100% 1,1,1,3,3,3-hexafluoro-2-propanol (TCI) at 1 mg/mL and incubated for 2 h at room temperature and then sonicated for 30 min. Subsequently, the solution was centrifuged 15000 × *g* and 4 °C for 30 min to remove the existing Aβ aggregates. The supernatant was collected and the solvent was removed by lyophilization overnight. The dried Aβ peptide was stored at −20 °C until needed.

### High-performance liquid chromatography (HPLC)

A Shimadzu LC-20AD system (Kyoto, Japan) was used for analytical HPLC separations. Samples were separated on a Shim-pack GWS C18 column (5 μm; 200 × 4.6 mm i.d.; Shimadzu GLC, Tokyo, Japan), and linear gradients were run from 0% B to 45% B over 45 min using 0.05% trifluoroacetic acid (TFA) in water (solvent A) and 0.05% TFA in acetonitrile (solvent B). Elution was monitored by measuring the absorbance at 536 nm. The flow rate was 0.5 mL min^-1^ and the column temperature 25 °C.

### UV–Vis spectroscopy

A UV-2450 (Shimadzu) spectrophotometer was used for UV-Vis spectroscopy. The concentration of betalain pigments without dextrin was determined using the molar extinction coefficient of ε = 65,000 M^−1^ cm^−1^ at 536 nm for purified red-beet betalain pigments.^28,29^

### ThT fluorescence assay

The ThT fluorescence assay was performed using the SensoLyte Thioflavin T β-Amyloid Aggregation Kit (ANASPEC, Fremont, CA, USA). Aβ and ThT solutions at final concentrations of 50 μM and 200 μM, respectively, were used in the evaluation system according to the manufacturers’ protocol. Aβ40 dissolved in phosphate buffered saline (PBS, pH7.4) or 10 mM NaOH and Aβ42 dissolved in 10 mM NaOH were used. Aβ solutions with red-beet betalain extract at final concentrations of 6.5, 12.5, 25, and 50 μM were evaluated. Morin, a flavonoid with inhibitory acitivity against Aβ aggregation,^30^ was used as an inhibitor control at a final concentration of 50 μM. A water sample without red-beet betalain extract or morin was used as the positive control, and a sample without inhibitor and Aβ40 was used as the reference control. ThT fluorescence was monitored at 37 °C using a spectrofluorometer (VarioskanLUX; Thermo Fisher Scientific, Waltham, MA, USA) at Ex/Em = 440/484 nm. Reading were taken every 5 min and each sample was shaken for 15 s immediately prior to the reading. Fluorescence data were analyzed using Skanlt software (Thermo Fisher Scientific). The reported values are the average of four wells.

### Circular dichroic (CD) spectroscopy analysis

For CD spectra analysis, 40 μM Aβ40 incubated with or without 200 μM red-beet betalain pigments for 7 days and 20 μM Aβ42 incubated with or without 100 μM red-beet betalain pigments for 4 days. The 7 days incubation sample of Aβ42 could not be measured due to precipitation. These incubated samples were measured using a CD spectropolarimeter (J-820, JASCO, Tokyo, Japan) at room temperature. Far-UV (190–260 nm) was measured with a scan speed of 100 nm/min. Every sample was scanned three times, and spectra presented are shown as the average of these results.

### TEM observation of Aβ aggregates

Aβ40 or Aβ42 was solubilized in DMSO or 10 mM NaOH to give a 1 mM solution, respectively. Aβ solutions were prepared with and without the addition of 50 μM red-beet betalain pigments and diluted to 20 μM with PBS. Aβ mixtures were incubated at 37 °C for 5 days before TEM observations.

Approximately 2 μL of sample solution was placed on a 150-mesh copper grid covered with formvar. After 5 min the sample was soaked away, then the grid was stained with 2.0% (w/v) uranyl acetate solution. TEM images of Aβ aggregates were obtained using a Hitachi 7650 TEM (Hitachi Co., Ltd., Tokyo, Japan) with an acceleration voltage of 80 kV

### *Caenorhabditis elegans* maintenance and synchronization

*C. elegans* were maintained on nematode growth medium (NGM) plates seeded with *Escherichia coli* OP50 (OP50) as a food source at 15 °C unless otherwise indicated. The transgenic strain CL2006 [dvIs (unc-54::human β-amyloid 1–42; pRF4)] was obtained from the Caenorhabditis Genetics Center (CGC; MN, USA). In CL2006, the human Aβ42 protein is expressed intracellularly within the body wall muscle. The expression and subsequent aggregation of Aβ in the muscle result in progressive paralysis.^22^

For synchronization, the nematodes were cultured on fresh NGM plates for 2–3 generations without starvation. Young adult nematodes were collected and cleaned, then disintegrated using a lysis solution (0.6% sodium hypochlorite and 200 mM NaOH). After 12–14 h at 20 °C, the isolated eggs were hatched to obtain the synchronized L1 larvae.

### Paralysis assay

The synchronized L1 larvae were fed OP50 and grown to young adults at 20 °C; 5-fluorodeoxyuridine (0.5 mg/ml) was added to prevent progeny production. The resultant synchronized hermaphrodites were transferred to OP50-seeded NGM plates containing various dextrin or red-beet betalain pigments concentrations (20 nematodes per plate). Dextrin was used as a control because the red-beet betalain pigments used in this study contained dextrin. The nematodes were tested for paralysis by tapping their noses with a platinum wire. According to a previous report, nematodes that moved their noses in response but fail to move their bodies were paralyzed.^31^ Paralyzed worms were calculated by the Kaplan-Meier method, and the difference in proportion of paralyzed worms was tested for significance using the log-rank test. Statistical analyses were performed using statistical software (GraphPad Prism ver. 8.0.1; GraphPad Software, Inc., La Jolla, CA, USA). Differences that gave \**p* < 0.05 were defined as statistically significance.

### Animals and surgery

Twenty-five male C57BL/6J mice (six-week-old) were purchased from Charles River Laboratories Japan (Yokohama, Kanagawa, Japan) and fed a standard rodent diet, CE-2 (CLEA Japan, Tokyo, Japan), for six weeks. At 12 weeks of age, mice were divided into five groups of five based on their body weight. Red-beet betalain pigments were suspended in distilled water at 83.3 mg/mL and its oral gavage carried out using a feeding needle (2.7 μmol/kg body weight as red-beet betalain pigments). After 1, 2, 3 and 6 h, mice were euthanized by CO_2_ inhalation and dissected. For the sham group, oral gavage of distilled water was performed and mice were immediately euthanized by CO_2_ inhalation. After blood sampling from the postcaval vein, and brain tissues were collected and frozen by using liquid nitrogen. Separated plasma and tissues were stored at −80 °C until analysis. Experiments were conducted following the guidelines for the appropriate conduct of animal experiments set by the Scientific Council of Japan (2006). The Animal Experimentation Ethics Committee of Ishikawa Prefectural University (Ishikawa, Japan) approved the study protocol (31–14–34).

### Quantification of betalain in brain samples

The amount of red-beet betalain pigments within brain samples was evaluated at selected time intervals (1, 2, 3, 6 h) after a single oral administration of the pigments. Brains, plasma, and red blood cells (five per time point) were then isolated and homogenized in PBS. Red-beet betalain pigments were extracted from samples with chloroform:methanol, at a ratio of 2:1 by volume (1 g of tissue with three volumes of extraction mixture). The methanol phase from all samples at each time point was dried, resuspended in 1% (v/v) acetic acid in the water and analyzed by HPLC.

## RESULTS

### Pigment analysis of red-beet betalain pigments

HPLC was used to investigate the composition of the betalain pigments present in the red-beet betalain extract. The red-beet betalain pigments, betanin and isobetanin were found to be the major pigments present in the extract, and other betalain pigments were not ascertained (Fig. 1). The proportions of betanin and isobetanin in this extract were almost equal (Fig. 1).

**Figure 1.**
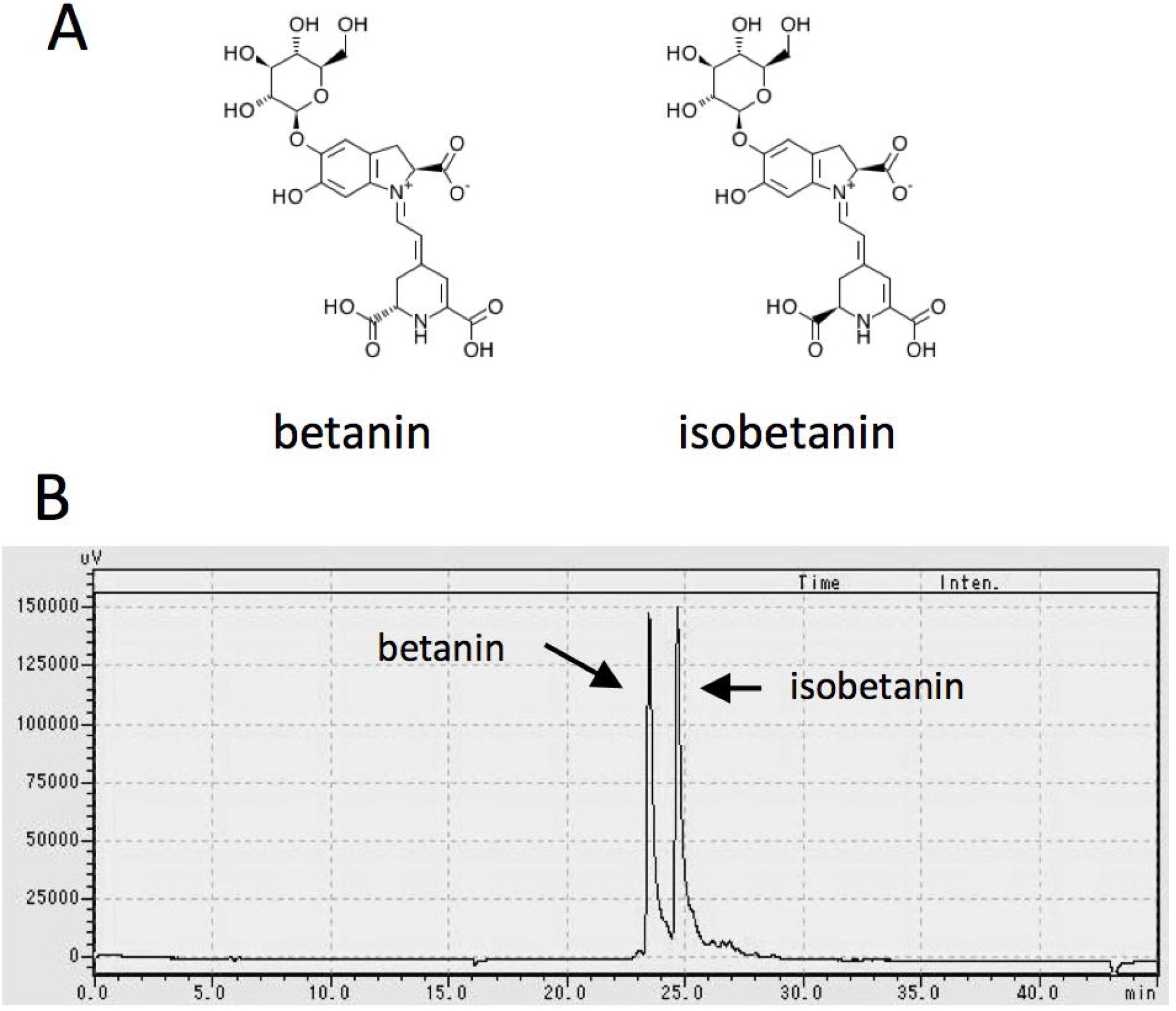
Composition of red-beet betalain pigments (A) Chemical structure of betanin and isobetanin (B) HPLC chromatogram of red-beet betalain pigments. HPLC elution sample peaks at 23.5 min and 24.6 min represent betanin and isobetanin, respectively.

### Red-beet betalain extracts inhibit the aggregation of Aβ40

Initially, the effect of red-beet betalain pigments on Aβ40 aggregation was investigated by using the ThT fluorescence assay. Thioflavin T used in this assay is a fluorescent dye that binds to amyloid and increases in fluorescence intensity. Aβ40 was dissolved in PBS, and red-beet betalain pigments at various concentrations (6.25, 12.5, 25 and 50 μM) were added. All samples with red-beet betalain pigments present had lower ThT fluorescence when compared with that of the Aβ40 alone sample (Fig. 2A), suggesting that red-beet betalain pigments inhibit formation of Aβ40 amyloids. Furthermore, the inhibitory effects were similar between 6.25 μM red-beet betalain pigments and 50 μM morin, an amyloid inhibitor (control) isolated from mulberry fruit (Fig. 2A). The red-beet betalains over 6.25 μM showed higher Aβ40 aggregation inhibitory activity than 50 μM morin. Moreover, the ThT assay was performed using Aβ42 and Aβ40 dissolved in 10 mM NaOH because Aβ42 hardly dissolved in PBS. The results showed that the fluorescence of these Aβ samples containing red-beet betalains pigments were lower than that of Aβ42 and Aβ40 alone, but higher than Aβ samples containing morin (Fig. 2B, C).

**Figure 2.**
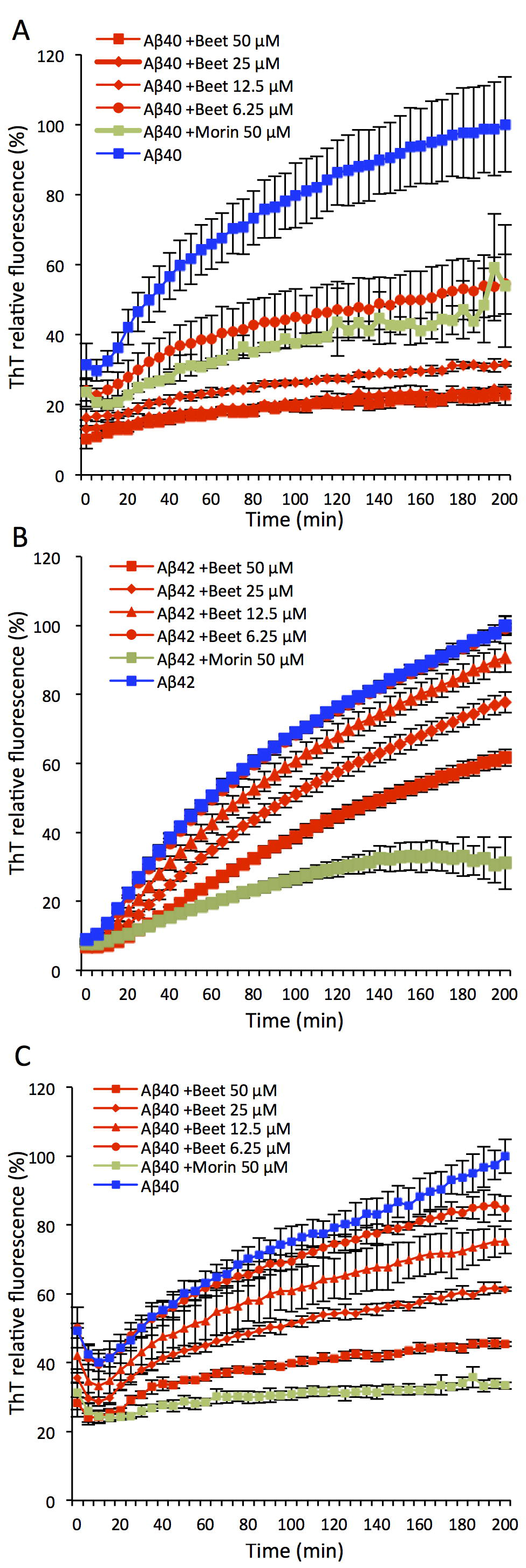
ThT fluorescence during amyloid fibrillation of human amyloid-β (Aβ). (A) Thioflavin T (ThT) fluorescence changes for Aβ40 incubated with/without red-beet betalain pigments. Aβ40 was dissolved in PBS. (B) ThT fluorescence changes for Aβ42 incubated with/without red-beet betalain pigments. Aβ42 was dissolved in 10 mM NaOH. (C) ThT fluorescence changes for Aβ40 incubated with/without red-beet betalain pigments. Aβ40 was dissolved in 10 mM NaOH. Aβ was incubated with 6.25, 12.5, 25 and 50 μM red-beet betalain pigments at 37 °C for 200 min. Error bars represent the means ± SD (*n* = 4). ThT relative fluorescence was expressed as a percentage of fluorescence for the Aβ alone sample, which had a maximum value of 100%. “Beet” and “Morin” represent sample with red-beet betalain pigments and inhibitor control, respectively.

### The effect of red-beet betalain pigments on the secondary structure of Aβ

CD spectroscopy was performed to detect secondary structure changes of Aβ in the presence of red-beet betalain pigments (a 5:1 molar ratio of the pigment to Aβ, Fig. 3). During the initial stage of Aβ aggregation, the CD spectra of Aβ40 and Aβ42 without red-beet betalain pigments showed a negative minimum at 196–199 nm, demonstrating that Aβ40 and Aβ42 adopted no secondary structure, i.e., random coil (Fig. 3A, C). After incubating at room temperature, the CD spectrum of Aβ without red-beet betalain pigments did not show a negative minimum at 196–199 nm but a new characteristic peak was appeared at 217 nm (Fig. 3B and D). These CD spectra showed that Aβ40 and Aβ42 underwent a structural transition from random coil to the β-sheet secondary structure.^32^ After incubation at room temperature, the CD signals at 217 nm arising from Aβ40 and Aβ42 in the presence of red-beet betalain pigments were significantly weaker when compared with the spectra of Aβ alone (Fig. 3B and D). This indicates that the amount of β-sheet structure in Aβ was clearly reduced by red-beet betalain pigments. These results suggested that red-beet betalain pigments hamper the onset of the conformational transition of Aβ40 and Aβ42 from random coil toβ-sheet.

**Figure 3.**
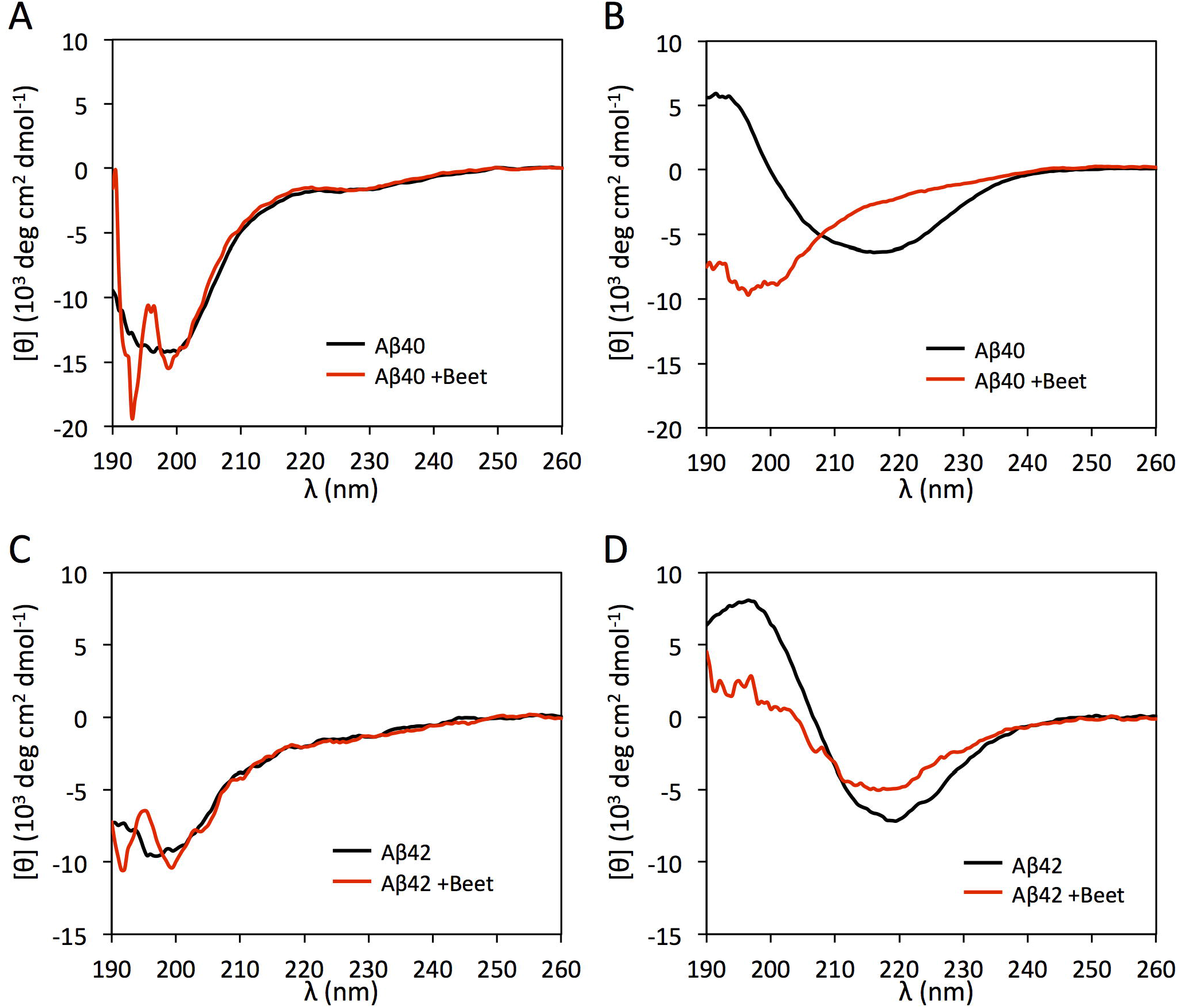
CD spectra of Aβ incubated with or without red-beet betalain pigment. (A) 40 μM Aβ40 incubated with or without 200 μM red-beet betalain pigment at 0 h. (B) 40 μM Aβ40 incubated with or without 200 μM red-beet betalain pigment at day 7. (C) 20 μM Aβ42 incubated with or without 100 μM red-beet betalain pigment at 0 h. (D) 20 μM Aβ42 incubated with or without 100 μM red-beet betalain pigment at day 4. Beet indicates red-beet betalain pigments.

### Red-beet betalain extracts modulate the morphology of Aβ Aggregates

The inhibitory effect of red-beet betalain pigments on Aβ aggregation was observed using TEM (Fig 4). Aβ40 and Aβ42 solutions (25 μM) with and without red-beet betalain pigments (50 μM) were incubated at 37 °C for 5 days, after which Aβ aggregation was monitored by TEM. Aβ fibrosis and amorphous aggregation were observed in Aβ40 and Aβ42 samples without red-beet betalain pigments (Fig. 4A, C). In contrast, Aβ fibisis and amorphous aggregation were significantly reduced in the sample with red-beet betalain pigments (Fig. 4B, D).

**Figure 4.**
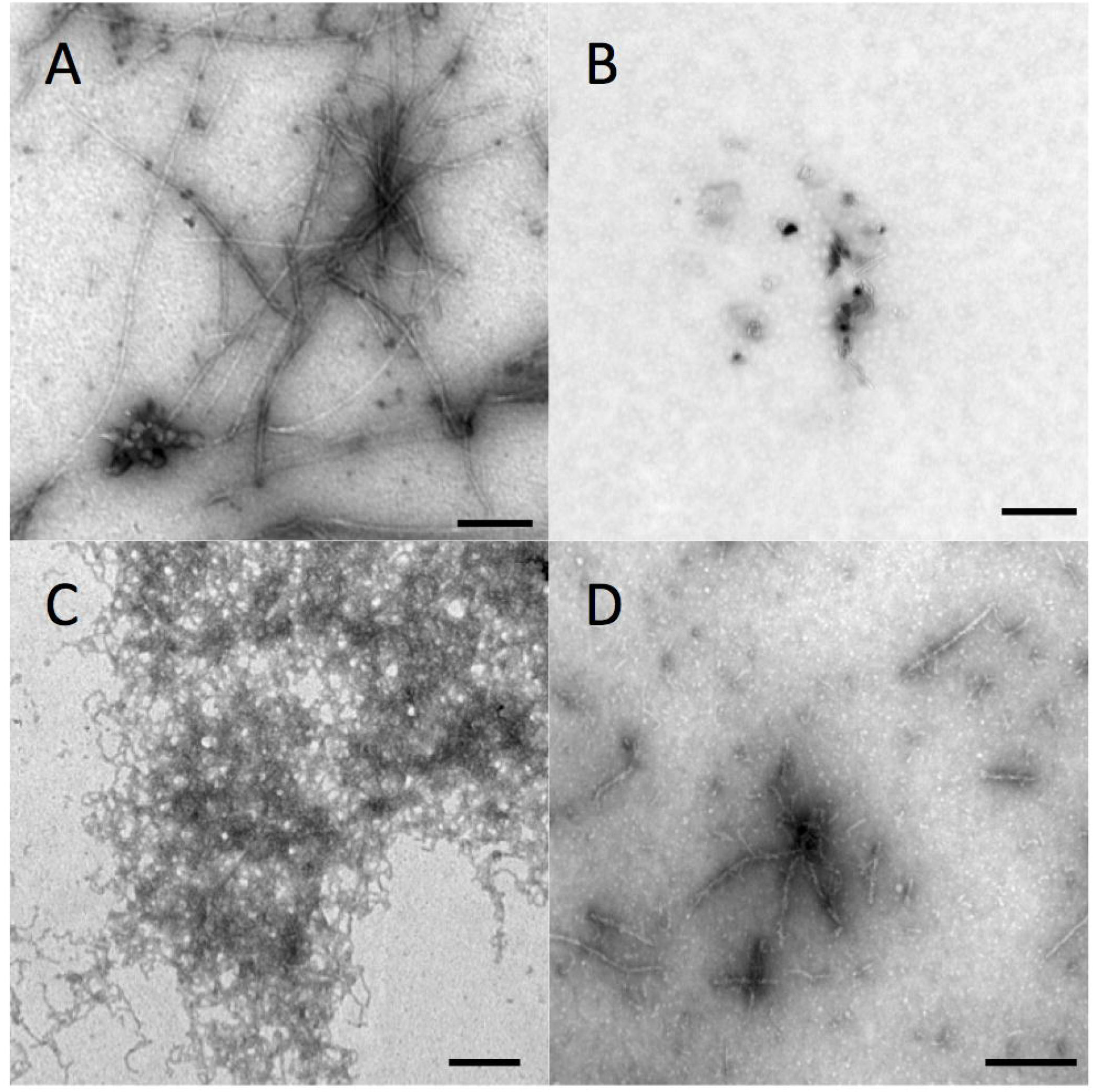
Transmission electron microscope images of human amyloid-β aggregates. (A) Aβ40 alone; (B) Aβ40 with 50 μM red-beet betalain pigments; (C) Aβ42 alone; (D) Aβ42 with 50 μM red-beet betalain pigments; Scale bars, 200 μm; Incubation time, 5 days.

### Red-beet betalain pigments delayed Aβ-induced paralysis in a transgenic nematode

*C. elegans* CL2006 strain expressing human Aβ in muscle develops a paralysis phenotype during grows. This transgenic nematode accumulates Aβ42, a major component of the senile plaques observed in AD, and is considered a nematode model of AD. We evaluated the inhibition of Aβ aggregation by red-beet betalain pigments *in vivo* using CL2006 nematode, as indicated in the experimental schedule presented in Fig. 5A. The nematodes were treated with dextrin or red-beet betalain pigments at concentrations of 2, 10 and 50 μM. As shown in Kaplan-Meier plots, dextrin added as a stabilizing agent for red-beet betalain pigments did not change the phenotype of the transgenic nematodes (Fig. 5B). In contrast, the highest concentration of red-beet betalain pigments (50 μM) delayed Aβ-induced paralysis in the transgenic nematodes significantly when compared with that of control (no betalain pigments), whereas the 2 and 10 μM treatments did not lead to a change in phenotype when compared with that of the control (Fig. 5C). Representative microscopic images treated (i.e., 50 μM red-beet betalain pigments) and control groups at day 15 are shown in Fig. 5D. By microscopic observation, many individual nematodes in the control group adopted a linear paralyzed body phenotype, whereas linear paralysis was not as distinct for nematodes in the group treated with 50 μM red-beet betalain pigments.

**Figure 5.**
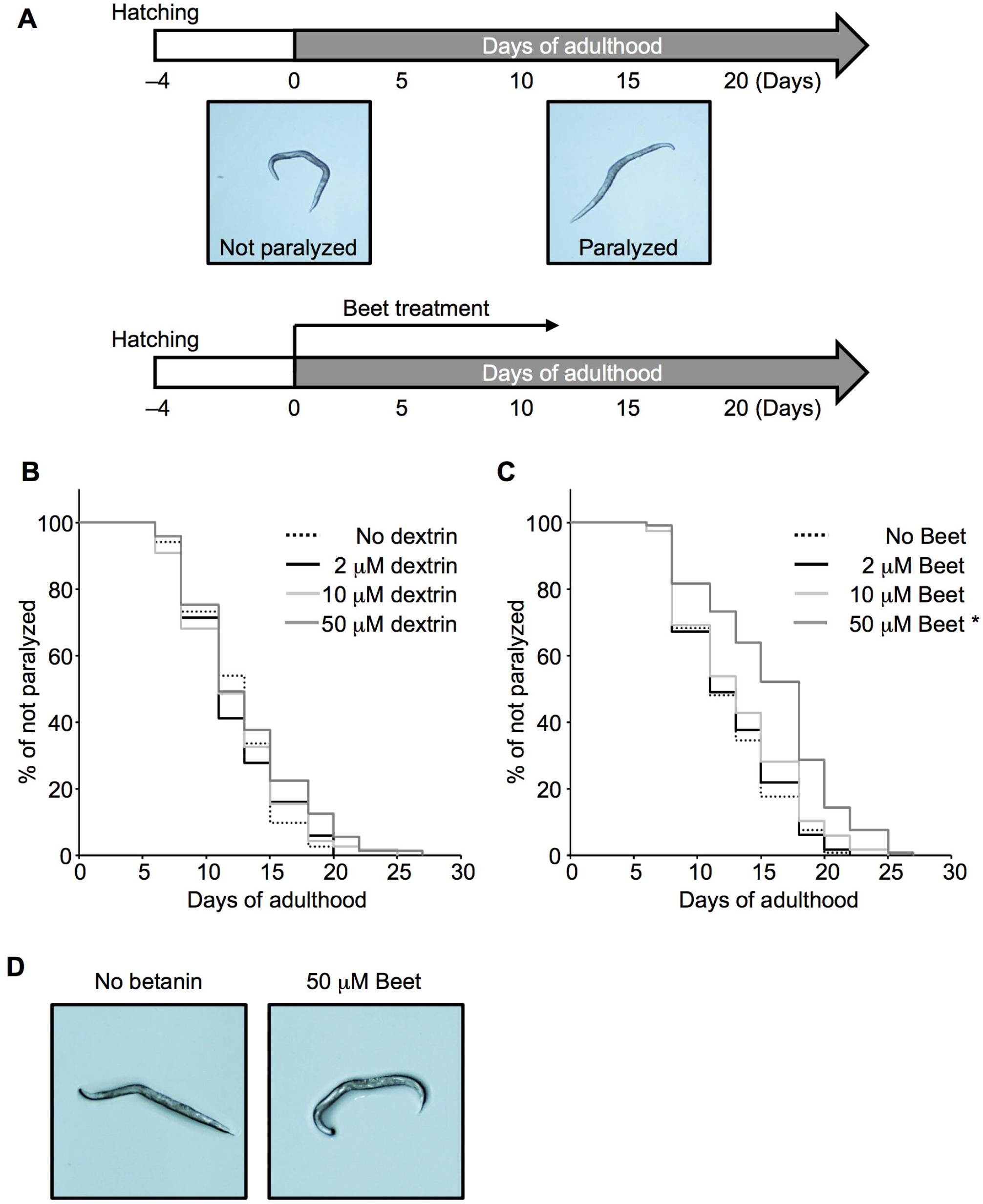
Effect of red-beet betalain pigments on amyloid-β-induced paralysis of *C. elegans* transgenic strain CL2006. (A) Experimental schedule of red-beet betalain pigments treatments in *C. elegans*. Image of *C. elegans* transgenic strain CL2006 morphology taken with a light microscope on day 0 (left) and day 15 (right) of adulthood. (B, C) The synchronized hermaphrodites were exposed to 2, 10, and 50 μM of dextrin (B) and red-beet betalain pigments (C). The paralysis rate was scored three times per week and is expressed as the percentage of not-paralyzed nematodes. (*n* = 109–119 nematodes/group). Differences in paralysis rates were tested for significance using the log-rank test (\**p* < 0.05). Results are representative of two independent experiments. (D) Morphological assessment of transgenic nematodes in the presence and absence of red-beet betalain pigments using a light microscope. Beet indicate red-beet betalain pigments.

## DISCUSSION

In this study, results from the ThT fluorescence assay, CD spectroscopy and TEM showed that red-beet betalain pigments composed of betanin and isobetanin inhibits Aβ aggregation. Furthermore, in feeding experiments using Aβ expressing nematodes, the ingestion of red-beet betalain pigments delayed the onset of paralytic symptoms caused by Aβ aggregation. Together, these results strongly suggest that betanin and isobetanin, which are the main components of red-beet betalain pigments, inhibit the aggregation of Aβ.

Previous studies have reported that betalain pigments have various biological effects, including anticancer,^10, 13^ anti-inflammatory^33^, inhibition of low-density lipoprotein oxidation^11, 12^ and inhibition of HIV-1 protease activity.^13^ Recently, nematode feeding experiments have shown that specific ingestion of betalain pigments produce lifespan-prolonging effects.^34, 35^ This study discovered an inhibitory effect on Aβ aggregation as a new physiological effect of betalain pigments.

PBS and 10 mM NaOH were used as solvents for Aβin the ThT assay. The results showed that the inhibitory activity of betalain pigments against Aβ aggregation was lower when the sample was dissolved in 10 mM NaOH versus dissolution in PBS. This observation indicated that pH affects the stability of betalain pigments. Betalains are reported to be stable between weakly acidic and neutral condition but are unstable outside this pH range.^36^ Therefore, the use of 10 mM NaOH may have destabilized the betalain pigments, which reduced their inhibitory activity against Aβ aggregation during the ThT assay. Nonetheless, red-beet betalain pigments are anticipated to be active against Aβ aggregation because the *in vivo* pH in human is near neutral.

Red-beet betalain pigments were found to suppress Aβ induced toxicity in transgenic nematodes. These transformed nematodes express Aβ42 in muscle but as a model organism do not have a blood-brain barrier (BBB). In mammals, red-beet betalain pigments must cross the BBB to accumulate in the brain and inhibit aggregation of Aβ. The betalain pigment, indicaxanthin, has been reported to cross the BBB in rats.^37^ As a preliminary experiment, we fed red-beet betalain pigments to mice and evaluated whether the pigments accumulated in the brain. However, betanin accumulation in the mouse brain was not confirmed (Suppl. Table 1). Nonetheless, a recent study reported that feeding betalain pigments to Alzheimer’s model rats alleviates AD symptoms.^38^ Thus, further *in vivo* studies are required to clarify whether betalains act directly to inhibit amyloid polymerization by passing across the BBB or function via a different mechanism such as modulating oxidative stress in blood.

## CONCLUSIONS

We found that beet-derived betalains inhibit the aggregation of Aβ. Additionally, feeding tests using Aβ-expressing *C. elegans* suggested that betalains inhibit Aβ aggregation *in vivo*. As a prospect, further analysis of betalain pigments may lead to their use as inhibitors of Aβ aggregation and as supplements to prevent AD. In this study, we conducted a bioactivity evaluation using *C. elegans*. The evaluation of phytochemicals bioactivity using nematodes is an effective initial selection process. There are many types of molecular species in betalain pigments.^39^ Future research efforts may lead to the discovery of betalain molecular species that function more effectively in inhibiting Aβ aggregation than red-beet betalain pigments. Furthermore, by using various model nematodes,we expect to find new bioactivities of betalain pigments.

## ACKNOWLEDGMENTS

The authors thank Akiko Mizuno, Hiroko Hayashi, Mami Awatani and Hitomi Nishikawa for their excellent technical assistance. This research did not receive any specific grant from funding agencies in the public, commercial, or not-for-profit sectors. We thank Edanz Group (https://en-author-services.edanz.com/ac) for editing a draft of this manuscript.

## CONFLICT OF INTERESTS

There is no conflict of interests to declare.

## CONTRIBUTIONS

MM conceived the study. TI and MM designed the experiments. TI performed the ThT fluorescence assay. HK performed TEM observations. YH conducted the nematode experiment. KM and YH conducted the experiment using mice. NI and SO performed CD spectroscopy analysis. TI, YH, KM, and MM wrote the draft manuscript. All authors have read and approved the final manuscript.

## SUPPORTING INFORMATION

Supporting information may be found in the online version of this article.

## Notes

### Competing Interest Statement

The authors have declared no competing interest.

### Summary of Updates

Aβ42 results were added to all *in vitro* experiments. Added the results of CD spectroscopy. Revised Fig2-5. Updated the author information.

